# Calibration of additional computational tools expands ClinGen recommendation options for variant classification with PP3/BP4 criteria

**DOI:** 10.1101/2024.09.17.611902

**Authors:** Timothy Bergquist, Sarah L. Stenton, Emily A.W. Nadeau, Alicia B. Byrne, Marc S. Greenblatt, Steven M. Harrison, Sean V. Tavtigian, Anne O’Donnell-Luria, Leslie G. Biesecker, Predrag Radivojac, Steven E. Brenner, Vikas Pejaver, ClinGen Sequence Variant Interpretation Working Group

## Abstract

**Purpose:** We previously developed an approach to calibrate computational tools for clinical variant classification, updating recommendations for the reliable use of variant impact predictors to provide evidence strength up to *Strong*. A new generation of tools using distinctive approaches have since been released, and these methods must be independently calibrated for clinical application.

**Method:** Using our local posterior probability-based calibration and our established data set of ClinVar pathogenic and benign variants, we determined the strength of evidence provided by three new tools (AlphaMissense, ESM1b, VARITY) and calibrated scores meeting each evidence strength. Results

All three tools reached the *Strong* level of evidence for variant pathogenicity and *Moderate* for benignity, though sometimes for few variants. Compared to previously recommended tools, these yielded at best only modest improvements in the tradeoffs of evidence strength and false positive predictions.

**Conclusion:** At calibrated thresholds, three new computational predictors provided evidence for variant pathogenicity at similar strength to the four previously recommended predictors (and comparable with functional assays for some variants). This calibration broadens the scope of computational tools for application in clinical variant classification. Their new approaches offer promise for future advancement of the field.

## INTRODUCTION

The classification of variants as pathogenic or benign by clinical genetic testing laboratories is a key component of modern genomic medicine. The American College of Medical Genetics and Genomics (ACMG) and the Association for Molecular Pathology (AMP) have made recommendations to standardize the practice of clinical variant classification.^1^ These recommendations identified distinct sources of evidence regarding the pathogenicity or benignity of a variant (e.g., genetic, functional, computational, case observation, and population data), assigned strengths to them, and specified rules to combine evidence to classify a variant into one of five classes: pathogenic, likely pathogenic, uncertain significance, likely benign or benign. Within the Richards *et al*. ACMG/AMP recommendations, the PP3 and BP4 criteria generally specified that evidence from computational tools (e.g., rule-based, statistical and/or machine learning-based) was considered to be the weakest, i.e., *Supporting* evidence. However, powerful, new variant impact predictors (VIPs) have rapidly emerged, with over 400 now developed.^2^

Recently, we undertook a rigorous quantitative calibration of computational tools, which demonstrates that some tools could reliably provide higher levels of evidence strength.^3^ Our approach maps scores from a computational tool to local posterior probabilities, which in turn, map to levels of evidential strength in the ACMG/AMP recommendations and their subsequent adaptation into a point-based system using a Bayesian formulation: *Indeterminate* or 0 points, *Supporting* or ±1 point, *Moderate* or ±2 points, *Strong* or ±4 points, and *Very Strong* or ±8 points.^4,5^ By applying this approach to 13 tools that predict the impact of missense variation, we demonstrated that at certain score thresholds, four tools can provide *Strong* evidence for pathogenicity and *Moderate* evidence for benignity: BayesDel,^6^ MutPred2,^7^ REVEL,^8^ and VEST4.^9^ Based on our findings, ClinGen^10^ recommended modifications to the PP3 and BP4 criteria that stipulated consistent use of a single tool defined in advance (per laboratory or per gene) with score thresholds calibrated to specific evidential strength levels up to *Moderate* benign (BP4_Moderate; -2 points) and *Strong* pathogenic (PP3_Strong; +4 points). Additional context about these clinical recommendations is provided in Stenton *et al*.,^11^ along with practical guidance on their intended use and their implications for variant curation in disease-associated genes.

Since then, advances in protein structure prediction, protein language models, and assay technologies such as deep mutational scanning (DMS) and massively parallel reporter assays (MPRAs), among others, have led to the emergence of new VIPs, with claimed improvements in predictive performance when compared to existing tools.^12–16^ However, it is unclear if these improvements in performance translate to the clinical context, in which computational tools serve as one line of evidence for variant pathogenicity/benignity among many. Furthermore, the objectives of these tools may vary, often focusing on the discovery of novel variants in research studies rather than the assertion of clinical pathogenicity, and predicting different notions of variant impact, e.g., distinguishing unobserved from observed ones. Thus, default score thresholds for these tools do not necessarily correspond to those for appropriate strength of evidence defined by the ACMG/AMP recommendations. Here, we estimate thresholds for newer tools corresponding to evidential strength in these recommendations, employing the same rigorous data sets and approaches. We also estimate additional thresholds for the above four previously calibrated tools corresponding to the ACMG/AMP point-based system for variant classification.^5^ We then compare and contrast these clinically performant methods against three recently published ones. Finally, we discuss our findings in light of the development and use of computational tools in the clinical classification of variants, reiterating the important role that we expect such tools to play in the future.

## MATERIALS AND METHODS

### Data sets, calibration procedures and *post hoc* analyses

We applied the methods and data sets developed in Pejaver *et al*.^3^ Specifically, we employed the *ClinVar 2019* data set for calibration and the *ClinVar 2020* set for *post hoc* assessments of tools and their thresholds. We used the *gnomAD* data set (v2.1.1) for both calibration and *post hoc* assessments.^17^ We calibrated each tool using our local posterior probability-based approach, and estimated score thresholds through bootstrapping, with the same parameters and local likelihood ratio cutoffs as before. We adopted the same *post hoc* assessment pipelines as in the Pejaver *et al*. study.

### Selection of computational tools and processing of their outputs

We selected tools for this study using a purposive sampling strategy. Based on recency of publication (within the past three years), the use of modern machine learning approaches (such as protein language models), their performance in the “Annotate All Missense” challenge^18^ in the Critical Assessment of Genome Interpretation (CAGI),^19^ anecdotal feedback on interest in adoption by the clinical genetics community, and the minimal need for access to original training data, we chose four tools for calibration: AlphaMissense,^15^ ESM1b,^14^ EVE,^12^ and VARITY^13^ (specifically, VARITY_R, the model trained on only rare variants). Important for this effort and also for utility within the clinical genetics community, these tools make precomputed scores for all possible single nucleotide or amino acid variants freely and publicly available, albeit in slightly different formats and with gene/protein identifiers from different databases.

We developed customized mapping protocols for each tool to maximize the number of variants in our data sets with scores. For AlphaMissense, we used three complementary mapping approaches. First, we linked precomputed scores to our data sets using chromosomal coordinates and Ensembl transcript identifiers as the key.^20^ Second, to ensure that the correct isoform was being considered, we undertook the mapping based on the Ensembl transcript identifier and amino acid substitution. Third, we undertook an additional mapping based on UniProt protein identifiers, using the corresponding mapping file provided by AlphaMissense.^21^ For ESM1b, we mapped precomputed scores to our data sets using the provided UniProt identifiers (with and without isoform-specificity) and amino acid substitutions. For variants that still remained unmapped, we used dbNSFP v4.4a^22^ to reannotate our variant list with the most up-to-date UniProt annotations, which were in turn used to map precomputed scores to our data sets. For EVE, we first mapped variants using UniProt or Ensembl transcript identifiers and amino acid substitution. We further matched all remaining unmapped variants to the UniProt gene name and amino acid substitution. For VARITY, we first mapped precomputed scores to variants in our data sets using UniProt protein identifiers, without consideration of the specific isoform. We then mapped the remaining variants strictly using chromosomal coordinates.

Except for VARITY, none of these tools were explicitly trained on variants from ClinVar.^23^ However, for VARITY, the precomputed score for each variant was assigned by a version of the model that did not include that variant in the training set. Therefore, no additional filtering of the data sets against the training data set of each tool was performed.

## RESULTS

### Recently published tools can provide up to *Strong* evidence for pathogenicity

Our local posterior probability-based calibration approach enabled the estimation of score thresholds for AlphaMissense, ESM1b, and VARITY_R that corresponded to distinct evidential strength levels within the ACMG/AMP variant classification guidelines. We found that all three tools were able to reach at least the *Moderate* level for benignity (with VARITY_R reaching *Strong*) (BP4) and the *Strong* level of evidence for pathogenicity (PP3) (Table 1, Fig. 1A). However, the score thresholds at which these were achieved were more stringent than the thresholds recommended by the tool developers. In fact, the recommended thresholds for AlphaMissense (0.564) and ESM1b (−7.5) do not meet the *Supporting* level of evidence for pathogenicity or benignity, based on our calibration. Overall, all three tools exhibited similar behavior to the four best-performing tools from our previous study, even when considering newer intervals between *Moderate* and *Strong* according to the ACMG/AMP point-based system (Table 1). When we attempted to calibrate EVE, it nominally appeared to reach the *Moderate* level of evidential strength for both pathogenicity and benignity. Score thresholds for *Supporting* and *Moderate* were 0.684 and 0.845, respectively, for pathogenicity, and 0.137 and 0.209, respectively, for benignity. However, EVE predictions were available only for a subset of genes in our calibration set, leaving about half of the benign/likely benign variants unscored. Furthermore, unscored genes showed a marked skew in ratio of pathogenic to benign variants. Due to potential sampling bias, we lack confidence in the applicability of the measured thresholds, rendering us currently unable to recommend their use in clinical variant classification.

**Table 1.**
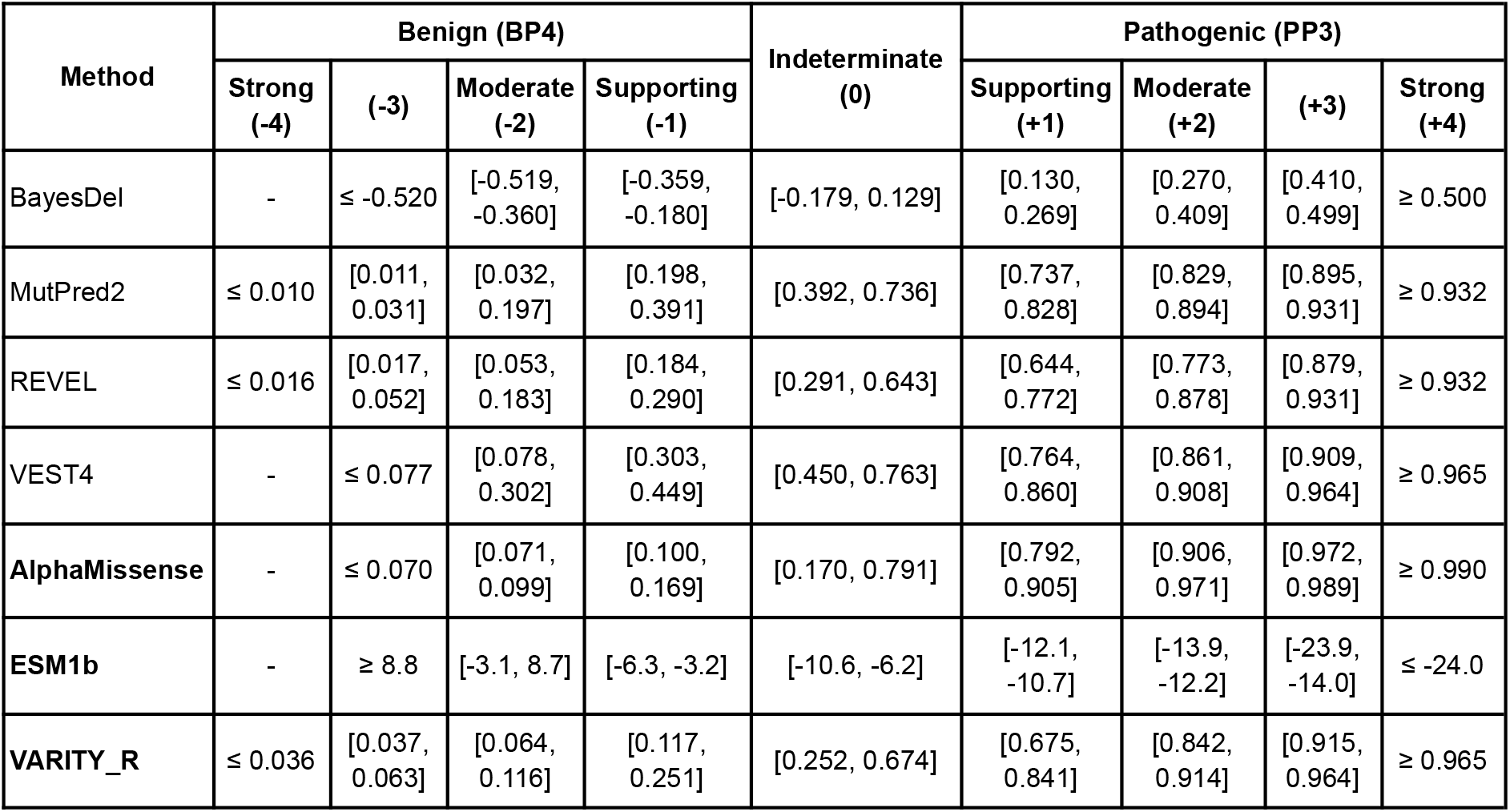
Estimated threshold intervals for all tools in this and our previous study according to the ACMG/AMP recommendations for sequence variant interpretation. The intervals correspond to the three pathogenic, one indeterminate, and three benign intervals (*Very Strong* not shown as it was never reached) in the current guidelines. The ACMG/AMP guidelines are expected to transition to a point-based system,^5^ and the numbers in parentheses in the header indicate point values corresponding to each evidential strength interval in this system. Although the 2015 guidelines do not include a strength level between *Moderate* (2 points) and *Strong* (4 points), intervals for the 3-point strength of evidence are also reported, as 3-point evidence will be recommended for future editions of the guidelines. A “–” implies that the given tool did not meet the posterior probability (likelihood ratio) threshold for that strength. All methods calibrated in this study are indicated in bold. For the remaining methods, all intervals are the same as those reported in our previous study,^3^ with additional columns for the interval corresponding to the Indeterminate range and ±3 points as per the point-based system.

| Method | Benign (BP4) |  |  |  | Indeterminate (0) | Pathogenic (PP3) |  |  |  |
| --- | --- | --- | --- | --- | --- | --- | --- | --- | --- |
|  | Strong (-4) | (-3) | Moderate (-2) | Supporting (-1) |  | Supporting (+1) | Moderate (+2) | (+3) | Strong (+4) |
| BayesDel | - | $\leq -0.520$ | [-0.519, -0.360] | [-0.359, -0.180] | [-0.179, 0.129] | [0.130, 0.269] | [0.270, 0.409] | [0.410, 0.499] | $\geq 0.500$ |
| MutPred2 | $\leq 0.010$ | [0.011, 0.031] | [0.032, 0.197] | [0.198, 0.391] | [0.392, 0.736] | [0.737, 0.828] | [0.829, 0.894] | [0.895, 0.931] | $\geq 0.932$ |
| REVEL | $\leq 0.016$ | [0.017, 0.052] | [0.053, 0.183] | [0.184, 0.290] | [0.291, 0.643] | [0.644, 0.772] | [0.773, 0.878] | [0.879, 0.931] | $\geq 0.932$ |
| VEST4 | - | $\leq 0.077$ | [0.078, 0.302] | [0.303, 0.449] | [0.450, 0.763] | [0.764, 0.860] | [0.861, 0.908] | [0.909, 0.964] | $\geq 0.965$ |
| **AlphaMissense** | - | $\leq 0.070$ | [0.071, 0.099] | [0.100, 0.169] | [0.170, 0.791] | [0.792, 0.905] | [0.906, 0.971] | [0.972, 0.989] | $\geq 0.990$ |
| **ESM1b** | - | $\geq 8.8$ | [-3.1, 8.7] | [-6.3, -3.2] | [-10.6, -6.2] | [-12.1, -10.7] | [-13.9, -12.2] | [-23.9, -14.0] | $\leq -24.0$ |
| **VARITY_R** | $\leq 0.036$ | [0.037, 0.063] | [0.064, 0.116] | [0.117, 0.251] | [0.252, 0.674] | [0.675, 0.841] | [0.842, 0.914] | [0.915, 0.964] | $\geq 0.965$ |

**Figure 1.**
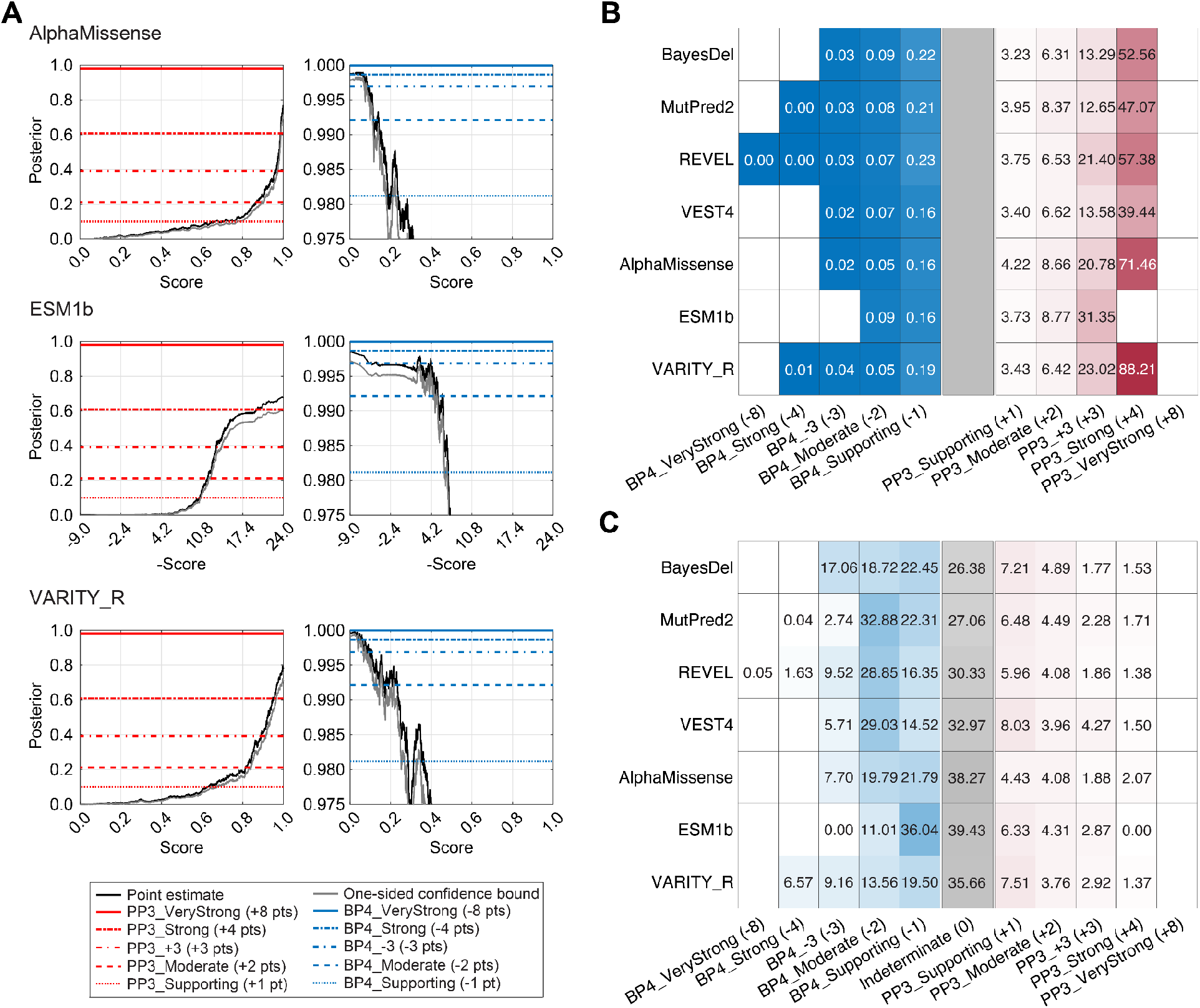
Local posterior probability curves and comparison with previously calibrated tools. **(A)** Pairs of curves for AlphaMissense, ESM1b and VARITY_R. For each tool, the curve on the left is for pathogenicity (red horizontal lines) and the curve on the right is for benignity (blue horizontal lines). The horizontal lines represent the posterior probability thresholds for *Supporting, Moderate, Strong*, and *Very strong* evidence as per current ACMG/AMP guidelines. A horizontal line representing the 3-point strength of evidence is also shown. The black curves represent the posterior probability estimated from the *ClinVar 2019* set. The gray curves represent one-sided 95% confidence intervals (in the direction of more stringent thresholds), calculated from 10,000 bootstrap samples of this data set. The points at which the gray curves intersect the horizontal lines represent the thresholds for the relevant intervals. **(B)** The likelihood ratios within each interval on the independent *ClinVar 2020* set. Darker colors indicate higher values for pathogenicity and lower values for benignity (because these are positive likelihood ratios). The limits for the color gradients are asymmetric, with ranges set between zero and one for benignity, and one and 100 for pathogenicity. A gray rectangle is introduced at the center for comparability with **(C). (C)** The percentage of variants predicted to be within the interval in the *gnomAD* set. Blue and red distinguish the evidential strength intervals for benignity from pathogenicity, respectively, with the indeterminate interval colored gray. The color gradient corresponds to the value in the cells, regardless of color. Darker colors indicate higher proportions. A white cell without a value indicates that the tool did not reach thresholds corresponding to that interval. The indeterminate interval also included variants without any scores.

### Clinical calibration shows modest improvements over existing computational predictors

We assessed the validity of our calibration by using the score thresholds estimated in Table 1 to group variants from the *ClinVar 2020* (not used in calibration) and *gnomAD* data sets, while also comparing them to the four previously calibrated tools (Fig. 1B and 1C). For the *ClinVar 2020* set, we calculated likelihood ratios within each interval defined by these thresholds, reflective of true and false positive rates for the classification of pathogenic variants. All tools met or exceeded (or, for benignity, were less than) the expected likelihood ratio values corresponding to each interval. The only exception to this was that some of the previously calibrated tools did not meet the thresholds for the 3-point intervals (Fig. 1B). VARITY_R and AlphaMissense resulted in higher likelihood ratios in the interval corresponding to *Strong* for PP3 than the four previously calibrated tools. However, it is unclear to what extent this is driven by the small number of variants in this interval relative to other intervals. No variant in the *ClinVar 2020* set received an ESM1b score of -24.0, effectively capping the maximal strength achieved by ESM1b at *Moderate* in practice. For the gnomAD set, we calculated the proportion of variants lying within each interval to assess how evidential strength is distributed for each tool in variants from the population (Fig. 1C). VARITY_R and AlphaMissense behaved as expected, in a manner similar to the four previously calibrated tools, with the proportion of variants in the *Strong* interval for pathogenicity being within the estimated prior probability of pathogenicity (0.0441). However, AlphaMissense classified the smallest proportion of variants as being within all three pathogenic intervals (0.125), slightly lower than REVEL (0.133). It is unclear if this results from AlphaMissense being trained on variants from gnomAD as a proxy for non-pathogenic variants.

## DISCUSSION

In this study, we calibrated three recently published computational tools to be usable within the ACMG/AMP guidelines for clinical variant classification and found that all tools reach evidential strength levels that are clinically useful. However, their recommended (default) thresholds did not meet even the *Supporting* level of evidence for variant pathogenicity. Furthermore, these three recent tools largely behaved similarly to four tools that we previously calibrated, and at best offer modest improvements in the strength of evidence that can be applied while minimizing the number of false positive predictions in the *Supporting* and *Moderate* categories. We also extended our previous study to include intervals corresponding to three points, in light of the point-based system to weight evidence that will be recommended in the next version of the ACMG/AMP standards. We did not calibrate methods that incorporate allele frequency as an explicit or strong implicit feature for two reasons. First, use of a predictor incorporating allele frequency will limit use of lines of evidence depending upon allele frequency, such as BA1, in variant classification. In practice, this means such methods are impractical to use in most clinical classification pipelines. Second, methods using allele frequency (AF) need to be calibrated distinctly for different AF thresholds (or once for the most stringent AF group), for which we currently lack sufficient data.^18^

This calibration shares the limitations of our previous study, including those related to the representativeness of data, potential circularities, estimation of prior probabilities, applicability and variability for specific genes and diseases.^3,24^ Of particular note, the gap in time between data set construction and the publication of some of these tools meant that there would invariably be irreconcilable differences among gene, protein and/or variant identifiers in our data sets compared to the files with precomputed scores for each tool. We expect this to be a major issue only if the differences in missing data were non-random, which was not the case here (average proportion of missing-at-random scores < 10%). For example, EVE^12^ was excluded because predictions were available only for a subset of genes in our calibration set, specifically leaving about half of the benign/likely benign variants in our data set unscored, and thus potentially introducing sampling bias.

The development of more advanced computational predictors of variant impact has often been motivated by the idea that no computational method can yet “be relied on alone for genetic diagnosis.”^25^ However, this is an inappropriate and unachievable benchmark for utility, because no single source of evidence other than high allele frequency–computational or otherwise–can presently be the sole criterion to determine the role of a variant in disease. Clinical standards for the classification of rare genetic variants always require the integration of multiple lines of evidence. This is a fundamental principle, integral to the ACMG/AMP clinical classification framework.^1^ As such, AlphaMissense authors’ assertion that it classifies “32% of all missense variants as likely pathogenic” employs the term “likely pathogenic” in a manner inconsistent with that used in clinical variant classification.

Historically, computational tools have been trained or calibrated to predict various proxies for variant pathogenicity that do not necessarily meet these clinical standards. As a consequence, their utility in clinical variant classification was initially limited to providing *Supporting* evidence. Our calibration provides a means to reconcile this misalignment of developers’ and clinical perspectives by providing data-driven, tool-specific guidance on use in clinical variant classification. We found that the AlphaMissense and ESM1b developers’ proposed thresholds did not achieve a *Supporting* degree of evidence, and our calibration recommends a higher threshold to reach *Supporting*. Our calibration also finds that for even higher thresholds, AlphaMissense and VARITY_R can reach *Moderate* and *Strong* pathogenicity evidence for some variants. This underscores the importance of independent calibration of methods used in clinical variant classification, just as critical assessments (such as CASP^26^ and CAGI^19^) have revealed how developers’ subtle knowledge of their methods and data inadvertently influence the results of their own assessments. Together with the ability to provide *Supporting* and *Moderate* benign evidence, we recommend these calibrated tools as potential alternatives alongside the previously recommended tools.

Our results continue to suggest increasingly important roles for computational predictors of variant impact in the interpretation of genomic data for clinical diagnosis and screening. The initial releases of this new generation of tools performed comparably to the best predecessors, suggesting potential for their future improvement. Moreover, the distinct approaches may offer independent information valuable for metapredictors. Relative to most other lines of evidence, computational tools have an outsized role because they can be readily applied to every relevant genomic variant. The continued development of enhanced in silico variant impact prediction methods augurs promising advances in clinical variant classification.

## DATA AVAILABILITY

Datasets described in the Materials and Methods are available on Zenodo: https://zenodo.org/records/13766399; intermediate result files and code to calculate local posterior probabilities, estimate thresholds, and plot figures in the paper are available here: https://github.com/pejaverlab/clingen-svi-comp_calibration. Machine-parsable calibration thresholds are available in VIPdb: https://genomeinterpretation.org/vipdb

## ACKNOWLEDGMENTS

This work was supported by NIH grants R00LM012992, R01HG013350 and U01HG012022. This work was also supported in part through the computational and data resources and staff expertise provided by Scientific Computing and Data at the Icahn School of Medicine at Mount Sinai and supported by the Clinical and Translational Science Awards (CTSA) grant UL1TR004419 from the National Center for Advancing Translational Sciences. Research reported in this publication was also supported by the Office of Research Infrastructure of the National Institutes of Health under award number S10OD026880 and S10OD030463. L.G.B. was supported by HG200388-10 and HG200387-10 and A.O’D.L. by U01HG011755. This work was conducted as part of the ClinGen Sequence Variant Interpretation Working Group. ClinGen is primarily funded by the National Human Genome Research Institute (NHGRI) with co-funding from the National Cancer Institute (NCI), through the following grants: U24HG009649 (to Baylor/Stanford), U24HG006834 (to Broad/Geisinger), and U24HG009650 (to UNC/Kaiser). The content is solely the responsibility of the authors and does not necessarily represent the official views of the National Institutes of Health.

## Author Information

Conceptualization: V.P., P.R., S.E.B.; Data Curation: T.B., V.P.; Formal Analysis: T.B., V.P.; Interpretation, discussion of results, and oversight: M.S.G., S.M.H., S.V.T., A.O.D-L., L.G.B., P.R., S.E.B., V.P.; Writing-original draft: T.B., V.P.; Critical evaluation of manuscript drafts, writing-review and editing: S.L.S., E.A.W.N., A.B.B., M.S.G., S.M.H., S.V.T., A.O.D-L., L.G.B., P.R., S.E.B., V.P.

## Ethics Declaration

This work does not report a clinical study or experiment with human subjects.

## MEMBERS OF MULTI-AUTHOR WORK GROUP

### ClinGen Sequence Variant Interpretation Working Group

Leslie G. Biesecker, Steven M. Harrison, Ahmad A. Tayoun, Jonathan S. Berg, Steven E. Brenner, Garry R. Cutting, Sian Ellard, Marc S. Greenblatt, Peter Kang, Izabela Karbassi, Rachel Karchin, Jessica Mester, Anne O’Donnell-Luria, Tina Pesaran, Sharon E. Plon, Heidi L. Rehm, Natasha T. Strande, Sean V. Tavtigian, and Scott Topper.

